# Fetal magnetoencephalography based on optically pumped magnetometers

**DOI:** 10.1101/2025.04.23.650121

**Authors:** Pierre Corvilain, Chiara Capparini, Vincent Wens, Lauréline Fourdin, Maxime Ferez, Odile Feys, Anne Delbaere, Rosine Lejeune, Deborah Goldman, Caroline De Coninck, Xavier De Tiège, Julie Bertels

## Abstract

The fetus in the third trimester of gestation has already the remarkable capacity to process external sensory information in utero. So far, investigations of fetal brain responses to sensory information have mostly relied on cryogenic magnetoencephalography (MEG), which is suitable to record fetal brain activity and is not much affected by layers of maternal tissues. Nevertheless, this solution is extremely expensive and limited to a couple of laboratories worldwide. In this work, we took advantage of the next generation cryogenic-free MEG, that is MEG based on optically pumped magnetometers (OPM), to develop a system that could record both fetal and newborn brain responses to auditory stimulation in a longitudinal design. Twenty-one pregnant women in their late third trimester of gestation (35-40 weeks of gestational age) were exposed to sequences of 500 Hz tones. Fetal brain activity was recorded using a wearable belt equipped with OPM sensors arranged on the women’s abdomen based on fetal head position. Results revealed that fetal OPM-MEG successfully recorded significant evoked brain responses to auditory stimuli that peaked ~300 ms post-stimulus at the group level. A similar auditory paradigm was performed with on-scalp OPM-MEG in 14 one-month-old infants, with 9 participants common to both timepoints. Infant responses showed a significant latency decrease compared to the fetal ones in terms of magnetometers; a decrease that did not reach significance level for virtual gradiometers. This work demonstrates the ability of OPM-MEG to non-invasively record fetal brain responses to external sensory stimuli. It paves the way for a wider use of fetal MEG to investigate fetal cognition and positions OPM-MEG as the most promising lifespan-compliant solution for monitoring early brain development.

## 1. Introduction

Early brain development has been increasingly studied over the past decades. Although developmental sciences have largely focused their investigations on the human infant, the origin of psychological development starts with fetal life (Reid & Dunn, 2021). Remarkably, brain architecture is already formed during pregnancy (Pletikos et al., 2014). In this context, previous research has shown that sensory systems are active even before birth. This was demonstrated by measuring fetal responsiveness (e.g., body and eye movements, heart rate variations) to stimuli coming from the extrauterine environment such as sounds (Hepper & Shahidullah, 1994; Visser et al., 1989) or lights (Kiuchi et al., 2000; Reid et al., 2017).

In line with these behavioral data, brain responses to external sensory stimulations have been demonstrated after 26 weeks of gestation (Vasung et al., 2019). While the interest in fetal brain imaging has increased over the years, its study remains challenging, with methods and analysis techniques that continue to emerge. Electroencephalography (EEG), which is the electrophysiological technique most widely adopted in infancy research (Azhari et al., 2020), is unfit for recording fetal brain activity. Indeed, bioelectric signals are strongly attenuated by several maternal tissue layers and the vernix caseosa coating the fetus from 25 weeks of gestation (Lengle et al., 2001). Investigations of the fetal brain have thus vastly relied on magnetic resonance imaging (MRI) and magnetoencephalography (MEG). MRI has been widely and reliably used to investigate the development of fetal brain structures (Manganaro et al., 2017; Saleem, 2014). Fetal brain function has also been investigated with resting-state functional MRI measurements, which allowed tracking brain networks maturation during pregnancy (Schöpf et al., 2012; Turk et al., 2019); see also (Zhang et al., 2019) for a review. However, task-based functional MRI investigations remain challenging, and only limited examples are available in the literature (Fulford et al., 2004; Jardri et al., 2008, 2012). These task-based studies are often characterized by a small sample size and restricted age range under investigation (Dunn et al., 2015). Furthermore, the reliance of fMRI signals on the neurovascular coupling (NVC) precludes the study of the fine-grained temporal and spectral dynamics of neural responses and their evolution with brain maturation. Finally, NVC is undergoing intense maturation that complicates the study of the hemodynamic responses linked to neural activity changes (Nourhashemi et al., 2019).

The most suitable non-invasive neuroimaging technique for investigating spontaneous and evoked fetal brain responses is fetal MEG (fMEG); see (Keune et al., 2019; Sheridan, Matuz, et al., 2010) for fMEG achievements and challenges. Accordingly, fMEG has been the most used imaging technique to investigate fetal brain activity over the past decades (Anderson & Thomason, 2013). Compared to the electrical potentials measured in EEG, the magnetic fields measured in MEG are less affected by the several layers of maternal tissues and muscles (Lowery et al., 2009), the skull (Lew et al., 2013) and the vernix caseosa separating the fetal brain from the sensors. Up to now, fMEG systems relied on superconducting quantum interference devices (SQUIDs), which require cryogenic cooling. The first fMEG study utilized a single SQUID to record fetal brain responses to auditory stimuli (Blum et al., 1985). More recently, multi-channel SQUID concave arrays, in which the whole maternal abdomen can fit in, have been designed (the SQUID Array for Reproductive Assessment, or SARA). The SARA system has undoubtedly increased the success in measuring functional neural activity in the fetus and allowed the characterization of both auditory (Draganova et al., 2005, 2007; Eswaran et al., 2021; Eswaran, Preissl, et al., 2002; Hartkopf et al., 2016) and visual (Eswaran et al., 2004, 2021; Eswaran, Wilson, et al., 2002; Matuz et al., 2012; McCubbin et al., 2007) evoked responses over the last trimester of gestation.

However, cryogenic MEG is an extremely expensive and rare solution to investigate the functional fetal brain. Fetal MEG is thus limited to a few laboratories worldwide (to the best of our knowledge, only two sites are equipped with cryogenic fMEG) and hence it remains unsuitable for large-scale fetal studies. Further, fitting a variety of abdominal shapes and sizes into a one-size-fits-all rigid sensor array poses some limitations, especially in terms of variable distance between the sensors and the abdominal surface (and in turn, the fetal brain), which is detrimental to the signal-to-noise ratio.

In this context, the next generation MEG, based on optically pumped magnetometers (OPMs), offers a cryogenic-free alternative and a promising solution for a wearable and more affordable fMEG system. Recent technological advances have made the measurement noise level of OPM-MEG comparable to SQUID-MEG (Boto et al., 2017), with OPMs showing better spatial resolution than SQUID arrays (Wens, 2023). Overall, the main advantages of OPM-MEG are the less constrained nature of data acquisition and the gain in brain-to-sensor distance. In fact, OPM sensors can be tailored to the users’ needs in wearable and adaptable configurations (Feys, Corvilain, Van Hecke, et al., 2023). Of note, they can be directly positioned onto the scalp or other surfaces of interest (as opposed to the vacuum space needed in SQUID-MEG) to reduce the brain-to-sensor distance, increasing the measurement sensitivity. Since the foundational work of (Boto et al., 2018), many studies have reported high-quality data with OPM-MEG in adults (Borna et al., 2020; de Lange et al., 2021; Seymour et al., 2021), in children (Boto et al., 2022; Feys et al., 2022; Hill et al., 2019; Rier et al., 2024) and even in newborns and infants (Corvilain et al., 2025; Feys, Corvilain, Bertels, et al., 2023).

OPMs have long been applied to fetal research in the context of monitoring cardiac activity in utero via magnetocardiography (MCG). Since the first successful fetal OPM-MCG recording (Wyllie et al., 2012), several studies have now used OPMs to create reliable, comfortable and cost-efficient fetal MCG systems (Escalona-Vargas et al., 2024; Eswaran et al., 2017; Strand et al., 2019; Wurm et al., 2023). These recent fetal OPM-MCG systems obtained results comparable to SQUID-based ones, at a fraction of the cost. Further, the portability of OPM sensors allowed to test pregnant participants in prone positioning, with improved comfort for the patient and higher signal amplitude compared to SQUIDs (Strand et al., 2019). Despite these achievements, measurement of fetal brain activity using OPM-MEG has until now remained elusive.

In the present work, we built an unprecedented OPM-MEG system to record fetal brain responses to auditory stimulation. To do so, we designed a practical and comfortable fabric belt covering the entire maternal abdomen with OPM holders directly sewn onto the fabric. The same OPM sensors were arranged onto an adjustable EEG-like cap setup for the neonatal phase of the research, as previously reported in (Corvilain et al., 2025). These setups were used to record auditory evoked responses to 500 Hz tones in a group of 21 fetuses at 35 to 40 weeks of gestation and a group of 14 neonates of 29 to 36 days of age (9 of the participants are common to both timepoints). Auditory evoked responses to 500 Hz tones have already been reported during the second trimester of gestation using SQUID-MEG (Eswaran et al., 2021; Eswaran, Preissl, et al., 2002; Sheridan, Draganova, et al., 2010), with detection rates in 50-90% of fetal subjects depending on gestational age and stimulus characteristics (Dunn et al., 2015). Past fMEG studies also provided information on the latency of these auditory responses, which decreased from the fetal to the neonatal period (Holst et al., 2005). We chose auditory stimulation since it has been associated with higher detection rates and more consistent fMEG findings compared to other sensory modalities (Dunn et al., 2015). We expect that fetal OPM-MEG will be more suitable not only for large-scale research studies, but also, in the future, for clinical investigations and management of at-risk pregnancies. This investigation is a further critical step towards making OPM-MEG a truly lifespan-compliant neuroimaging tool, capable of measuring brain activity from fetal to adult developmental stages.

## 2. Materials and Methods

### 2.1 Participants

Twenty-one pregnant women aged 27 to 39 years old (mean: 32 years) with uncomplicated pregnancies participated in this study. Data from 2 of them were rejected due to technical issues (abnormal noise floor). Of the remaining 19, 9 were primiparous, 8 had already given birth once, and 2 twice. Thirteen fetuses were female, with a gestational age between 35 and 40 weeks at the time of testing. All fetuses were in cephalic position. One fetus was the result of medically assisted procreation.

Nine women returned for the second timepoint of this longitudinal study when their child was between 29 and 36 days old (mean: 33 days), corresponding to an average of 45 days after their first visit (range: 35-55 days). Six of these infants were female. These nine newborns belonged to a bigger cohort that consisted of 14 full-term newborns (13 with usable data; 7 females), with ages ranging between 29 and 36 days (mean: 32.6 days). These neonatal data were analyzed in detail in (Corvilain et al., 2025). Detailed information about the fetal and newborn sample of the current study can be found in Table S1 in the Supplemental Material.

This study was approved by the Institutional Ethics Committee (P2021/141/B406202100081). The protocol was carried out in accordance with the approved guidelines and regulations. All participating mothers gave written informed consent prior to testing and received monetary compensation for their participation.

### 2.2 System design

We developed a fetal OPM-MEG system that is easy to set up and comfortable to use. The homemade system consisted of a stretchy maternity belt mounted with 45 to 54 (mean: 50, SD: 3) biaxial (mean: 29, SD: 2) and triaxial (mean: 21, SD: 2) rubidium-based OPMs (biaxial, Gen2 QZFM; triaxial, Gen3 QZFM, QuSpin Inc., size: 1.2 × 1.7 × 2.6 cm^3^, weight: 4.5–4.7 g), see Figure 1a-b. The OPMs were attached to the belt using custom OPM holders (3D-printed in acrylonitrile butadiene styrene), sewn onto the belt. As a result, the interior of the belt was all fabric, making it comfortable to wear for the pregnant participants. The holders were set so that a gap of 4 mm was left between the maternal skin and the OPM (see Figure 1c), which is crucial to dissipate the heat generated by the rubidium-based OPMs and prevent hurting the sensitive skin of the maternal abdomen. Indeed, the rubidium inside the sensor’s cell needs to be heated to approximately 150°C to work in the spin exchange relaxation-free regime (Tierney et al., 2019), making the OPM surface relatively hot. In addition, we attached a layer of aerogel foam to the bottom of the OPMs to further isolate the maternal skin.

**Figure 1.**
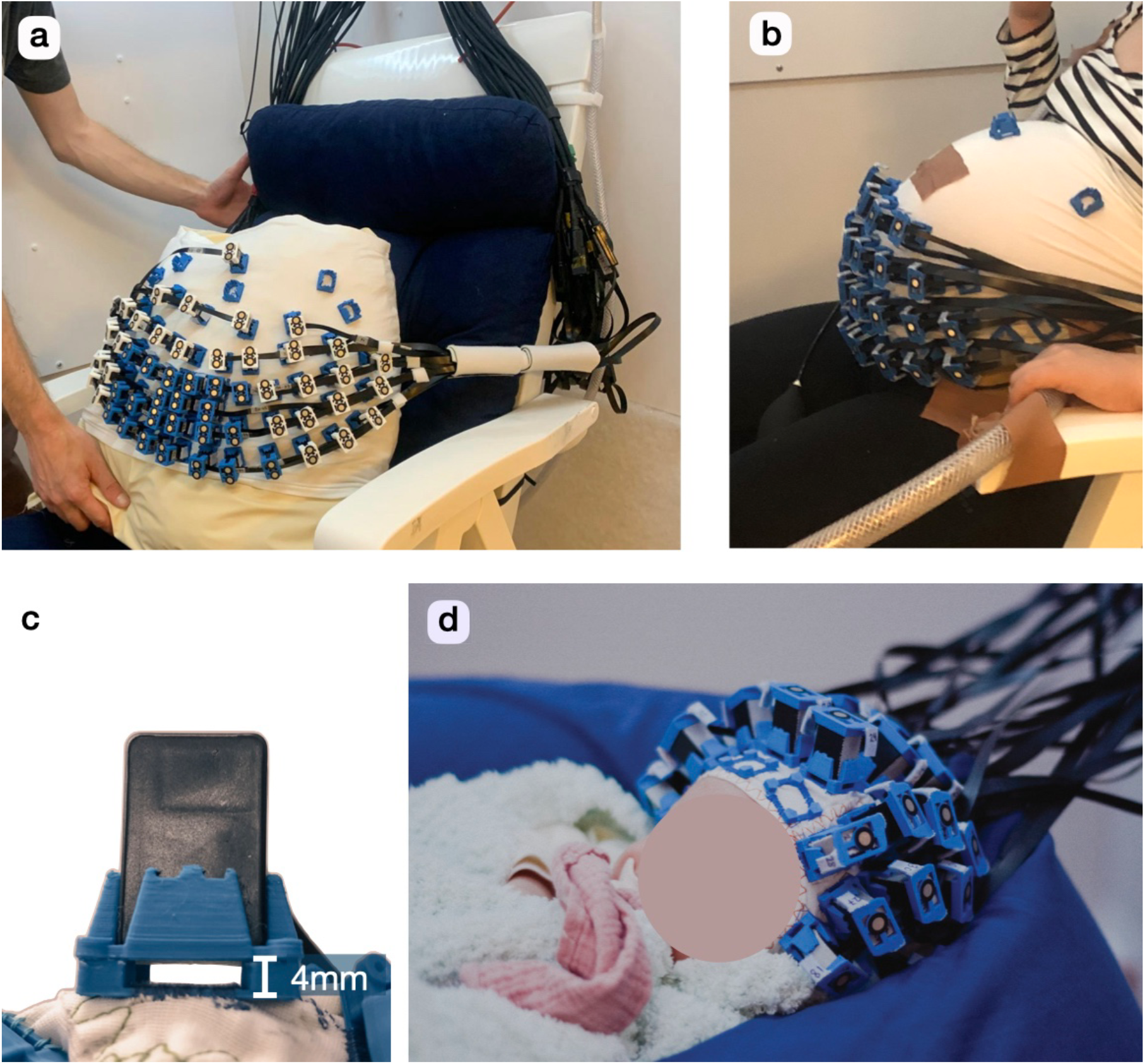
Fetal and neonatal OPM-MEG systems design. (a-b) Fetal OPM-MEG belt. (c) Small gap left between the OPMs and the abdomen or scalp skin for heat dissipation. (d) Neonatal OPM-MEG flexible cap.

The OPM density was increased in the central region of the belt (see Figure 1a), which was positioned right above the fetal head location (determined just before the recordings by ultrasonography). The triaxial sensors were prioritized in that area. Once the belt was installed, the pregnant women sat comfortably in an armchair throughout the experiment and watched a movie with the sound delivered through earplugs. A second ultrasound scan was carried out after the experiment to ensure that the position of the fetus did not change substantially. For the newborn timepoint of this research, we used the same OPM mounts sewn on flexible cap with similar OPMs as described above (see Figure 1d; for further details, see (Corvilain et al., 2025)).

### 2.3 Data acquisition

The experiment took place inside a magnetically shielded room (Compact MuRoom, MSL Ltd), and the remaining magnetic field background was further reduced below 1.5 nT using degaussianization and field nulling coils (MuCoils, Cerca Magnetics). In fetal OPM recordings, an accelerometer was placed on the maternal abdomen to record its movements. The OPM data, audio data and the accelerometer data were recorded simultaneously at a sampling frequency of 1200 Hz. Live video was available during the whole experiment to monitor and address any problems that may have occurred. For details of the field nulling and data recording procedures, see (Corvilain et al., 2025).

### 2.4 Auditory stimuli

The auditory paradigm consisted of 7-minute sequences of 500 Hz pure tones. Of note, high frequencies

(> 500 Hz) are reduced in utero by a combination of amniotic fluid inside the fetal ear and maternal tissues separating the ear and the sound source, whereas low frequencies (≤ 500 Hz) seem enhanced (Gerhardt & Abrams, 1996). For fetal OPM recordings, each tone lasted 100 ms including 15 ms of fade-in and fade-out, with a stimulus onset asynchrony (SOA) varying between 500 and 900 ms (500 to 700 ms for n=13 and 700 to 900 ms for n=6, both uniformly distributed). The tones were presented at 95 dB using a flat MEG-compatible speaker (Panphonics), which corresponds, after sound attenuation by the maternal tissue, to roughly 60-70 dB for the fetus (Querleu et al., 1988; Philbin, 2017). For neonates, SOAs were shorter on average (500-600 ms) and the sounds were presented at ~60 dB. The sequences were generated using a custom-made Python program (Python Language Reference, version 3.7). With four fetuses, we presented the sequence a second time. The mean recording time for this study was 8.6 ± 3 min. The number of tones presented per fetus ranged from 526 to 1122 (mean: 799, SD: 161); for the newborns, it ranged from 547 to 1751 (mean: 1038, SD: 412).

### 2.5 Data analysis

The data analysis was performed with custom-made scripts for Matlab (MathWork Inc., version 2020a-2024b), unless otherwise stated.

#### 2.5.1 Decomposition into independent components

As a preliminary validation of our setup, we used the fetal OPM recordings to extract MCG activities. Recording periods containing large movement artifacts were first identified by visual inspection of the continuous data and marked as artifact time windows. Data were bandpass filtered between 1 and 40 Hz. An independent component analysis (ICA) (Vigário et al., 2000) was applied to the artifact-free part of the recordings, using the FastICA algorithm (Hyvarinen, 1999). Components containing the heartbeat signals from the fetus and the mother were visually identified.

#### 2.5.2 Heart rate analysis

The heart components were initially classified into maternal or fetal by visual inspection, based on the expectation that R-peaks rates range from 60 to 110 bpm for maternal and 110 to 160 bpm for fetal cardiac activity (Ayres-de-Campos et al., 2015). Each component was then converted into a time course by computing heart rate as the inverse of the time difference between 2 successive R-peaks, assigning this value at the middle time point between the two R-peaks, and interpolating. R-peaks were extracted automatically by scanning the component signal within sliding windows (fetal: 5 s, maternal: 10 s; corresponding to roughly 10 beats per window) and identifying peak time points exceeding 3 SD within the window. If no R-peak was detected (e.g., due to non-cardiac interferences in the component), the threshold was lowered to 2SD. If there were several components, the corresponding heart rate time courses were averaged. Before and after averaging, time points where the interpolated heart rate derivative exceeded 2SD were discarded to eliminate inconsistent R-peak detections. Heart rate variability was assessed as the standard deviation of the heart rate time course across the 7 min sequences. For the remaining analyses focused on brain activity, heart components were removed from the fetal and neonatal OPM data.

#### 2.5.3 Virtual Gradiometers

Despite substantial steps for artifact suppression described below, fetal OPM signals remained rather noisy. When a magnetometer array is dense enough, a possible analysis method to remove part of the noise is to convert the signal to virtual gradiometers. Indeed, gradiometers are essentially capturing the difference in the magnetic field between two close-by points and are therefore insensitive to the homogenous (i.e. spatially constant) part of the noise (Hari & Puce, 2023). This technique has been fruitfully applied in the past (Hill et al., 2019). In our fetal OPM-MEG setup, the OPMs of interest were very dense, enabling us to convert magnetometers into virtual gradiometers. For each layout position, 6 virtual gradiometer signals were computed, one for each of the 2 directions tangential to the abdomen, times the 3 magnetic field components. They were computed as the mean of the difference between the magnetic field component measured at that position and at its nearest neighbors, projected onto the tangential direction in question. The nearest neighbors were determined as the ones that were not more distant than 1.58 times the distance to the first neighbor. If using that procedure less than 5 neighbors were found, the threshold was extended to 1.9. The step of computing the virtual gradiometers was performed right after removing the heart components. All the other steps described below were applied to the virtual gradiometers data as well.

#### 2.5.4 Artifact removal

Time windows affected by remaining artifacts were identified by visual examination (5 s sliding windows) of the non-heart independent components of the magnetometers data. The amount of data identified as contaminated by artifacts is described in Table S2 in the Supplemental Material. To avoid filtering artifacts, data in the artifact periods were replaced by linear interpolation before being bandpass filtered between 0.15-20 Hz and notch filtered at the frequencies at which the power spectrum displayed noise peaks, within the 13-20 Hz range. Artifacts periods were extended by 1.5 s in both directions. At this point, two specific steps were taken for noise suppression in fetal recordings. First, large signal artifacts associated with the mother’s respiration were removed using a non-stationary regression based on accelerometer data. More specifically, for each period longer than 4 seconds not defined as an artifact, we linearly regressed the first principal component of the 3-directional acceleration signal from the OPM data. Second, we defined 32 sensors of interest at the lowest and most central positions of the belt, which were most likely to measure cerebral signal based on the positions of all the fetal participants’ heads. The data from the other sensors were used as external noise reference and thus linearly regressed from sensors of interest using non-stationary multiple regression, again in artifact-free windows longer than 4 seconds. Artifact-free windows smaller than 4 seconds were rejected. Then, both restricted fetal and neonate data were high pass filtered at 1 Hz. A robust z-score rejection (channel per channel, threshold value of 5, data amplitude smoothed with a 1 s square window beforehand) was performed to identify samples affected by remaining artifacts. Finally, the first three principal components were projected out (Feys et al., 2022).

#### 2.5.5 Evoked response analysis

The resulting data were decomposed into epochs from −100 to 700 ms around the start of each sound presentation (trial), and epochs that intersected artifact periods were rejected. The mean number of trials kept was 550 (SD: 172) for the fetal timepoint and 694 (SD: 328) for the neonatal timepoint. The number of recorded and accepted trials per fetus participant is available in Table S2. The remaining epochs were baseline corrected, with a baseline spanning the 100 ms pre-stimulus period, and averaged for each participant, yielding individual-level evoked responses. At this stage, the responses were still contaminated by a stimulation artifact, which occurred at the same time as the sound stimuli (from 0 to 100 ms) and had the shape of the sound envelope. It most likely originated in the high sound pressure making the OPMs vibrate, resulting in an OPM signal shift at low frequencies. Some sensors and participants were more affected by this artifact than others. To deal with it, we computed the correlation of each signal with the sound envelope, which resulted in a spatial map of how the sensors were affected by the artifact. That map was spatially regressed, resulting in artifact-free responses. Finally, noisy channels were determined as those with a temporal SD during baseline exceeding the average across all channels by 2SD (separately for magnetometers and virtual gradiometers), and they were discarded (number of noisy channels: magnetometers, range: 1-7, mean: 4, SD: 1.4; virtual gradiometers, range: 5-10, mean: 8, SD: 1.5). Topographic plots of the resulting responses were generated using Fieldtrip (Oostenveld et al., 2011) based on a custom-made 2D template layout that most closely matched the sensors positions on the belt or cap. To obtain a group-level response in fetal data, one cannot simply average the responses channel per channel, as the variations in the position and orientation of the fetal heads and of the OPMs (variations in the belt positioning and abdomen morphologies) lead to individual topographic maps that vary substantially (see Figure S1). The approach taken here was to extract a single curve per participant that captures the temporal dynamics of the response and discards the spatial distribution, an approach adapted from (Moser et al., 2019). This response curve was computed separately for magnetometers and virtual gradiometers by first extracting the 5 channels contributing the most to the response, based on their weight in the first principal component of the response. Then the first principal component (PCA1) of those 5 channels was taken and its scale was set to match that of the root-mean-square of those 5 channels. These single curves per participant were taken to capture the response. Of note, this procedure was also performed on neonatal data even though standard response averaging could be done in this case (Corvilain et al., 2025). The latencies and amplitudes of the individual responses, and the group-average response were obtained from these curves. To avoid picking up noise peaks, the latencies were derived from a polynomial fit of PCA1 curves, whose degree was determined to minimize the latencies variance for each group (fetal: 8, neonatal: 15), as noise peaks increase this variance. In principle, response latency should not depend on whether it is based on magnetometers or virtual gradiometers, so the subjects for which the latency difference between the two sensor types was too large (greater than the mean plus 1.5 SD across participants) were discarded for the group analysis (fetal: n=4, neonatal: n=2). As the sign of the PCA1s are random, group-level response curves were obtained by averaging individual curves after a sign-flip ensuring positivity at the group-average latency. To further characterize the quality of the individual responses, we computed their lagged correlation with the group-average response, allowing shifts in latency up to 1.5 SD of all the latencies and taking the maximum lagged correlation.

### 2.5.6 Statistics

The fetal and maternal heart rate variability were compared by performing a two-tailed paired sample t-test (n=19), after carrying out a Shapiro-Wilk test that did not hint at a deviation from a normal distribution (W<1, *p*>0.05). Maximum statistics tests were performed on the PCA1 curves of the evoked responses at both the individual and group levels, hence providing full control of the familywise error rate over the period 100-700 ms post-stimulus (100-500 ms for the neonatal responses). The iterative step-down method (Nichols & Holmes, 2002) was used, i.e., time points that were found to be significant in one iteration were masked off the maximum statistics test in the next iteration until no more significant data were found. Null distributions were constructed from N=10,000 samples of surrogate data built according to the null hypothesis that they contain no response to the stimuli. Specifically, the sign of each epoch was randomly flipped (permutation test). These flips were applied to all the OPM channels (magnetometers and virtual gradiometers) at once to preserve their spatial distribution. Statistical tests were also applied to the correlations between the individual PCA1s and the group average. Null distributions (N=10,000) of correlations were constructed by randomly scrambling the Fourier coefficients of the group-average PCA1 curve before computing the correlation. Finally, latencies comparisons between the fetal and neonatal timepoints posed the challenge that half the data points were paired (subjects participating in both studies), and half were not (subjects participating in only one of the studies). We calculated a mixed *t*-statistics following (Yu et al., 2012) and generated a surrogate distribution (N=10,000) by randomly permuting the paired elements and shuffling all the unpaired ones. For all tests, the statistical significance level was set to *p* < 0.05.

## 3. Results

### 3.1 Fetal magnetocardiography

Performing an ICA on the fetal OPM-MEG data readily led to the identification of fetal and maternal heartbeats components (see Figure 2a). For all participants, we were able to extract the heart rate as a function of time from those magnetic field heart components (see Figure 2b). As a proof-of-concept for these fetal MCG measurements, we compared the fetuses’ heart rate variability to that of their mothers (Figure 2c); the difference was significant. Individual mean heart rate and heart rate variability values are reported in Table S1 in the Supplementary Material. The mean fetal heart rate was 140 bpm (SD: 9 bpm, range: 127-157 bpm).

**Figure 2.**
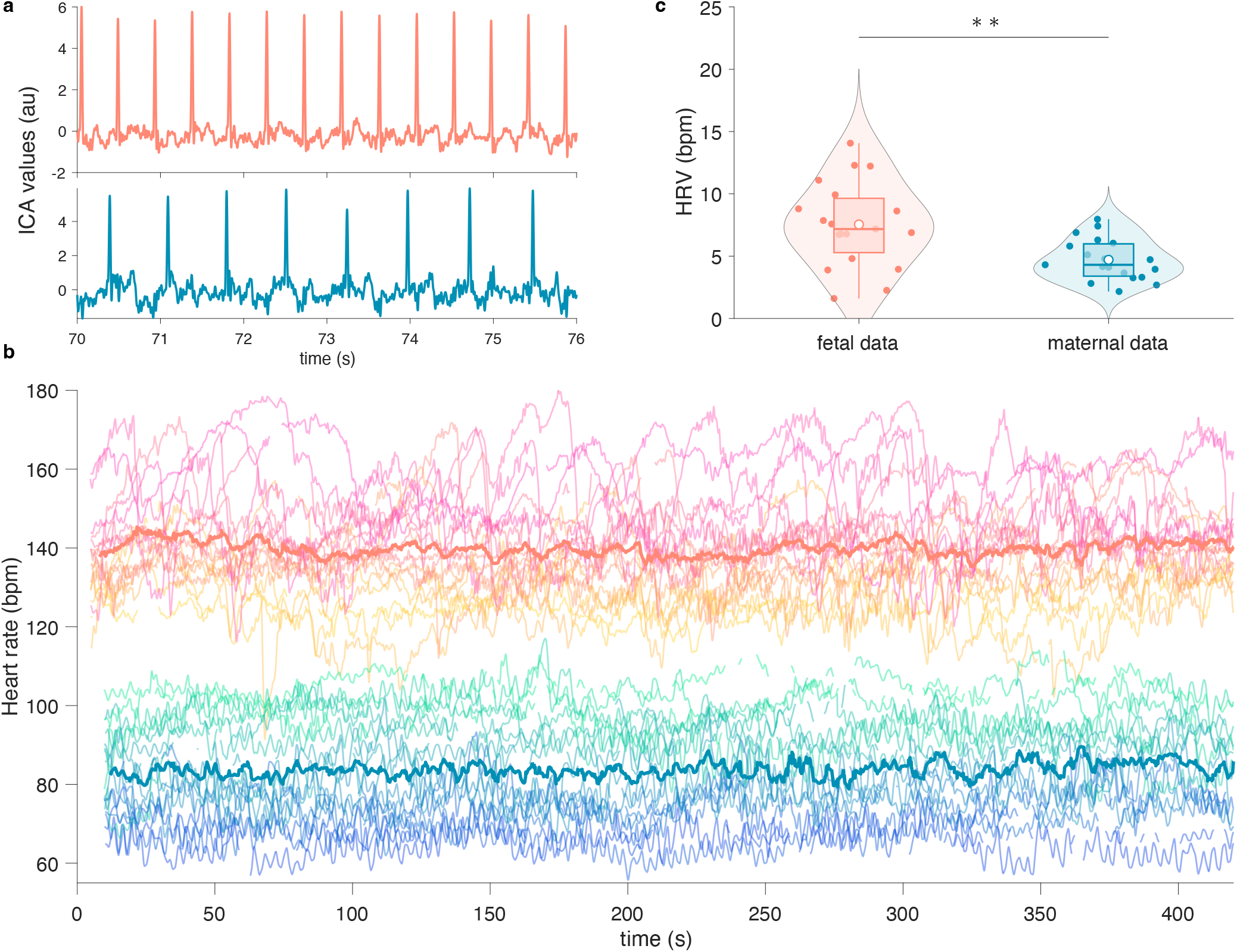
Fetal Magnetocardiography. (a) 6-s excerpts of the components corresponding to the fetus’ (top) and mother’s (bottom) heartbeats, for one participant. au = arbitrary units. (b) Individual heart rates in bpm as a function of time for all participants (if more than 7 minutes were recorded, only the first 7 minutes are included). The fetuses’ time series are in the top half (yellow to pink); the mothers’ ones in the bottom half (blue to green). The bold lines correspond to the group averages. bpm = beat per minute. (c) Comparisons of the heart rate variabilities (HRV) of the fetuses and that of their mothers. ** p<0.01.

### 3.2 Auditory evoked responses

#### 3.2.1 Fetal responses

Auditory evoked responses were evaluated at the individual level (Figure 3). Their PCA1s were extracted and are superimposed in red in Figure 3. In most fetuses, an evoked response was observable from 100-200 ms to 400-600 ms post-stimulus, with a peak at about 200-400 ms post-stimulus. Significant PCA1 responses were found in 5 fetuses. The group average of the PCA1s (excluding n=4 participants with discordance between magnetometers and virtual gradiometer latencies, see below) is displayed in Figure 4. The group response was significant (*p*<0.0001) from 279 to 363 ms post-stimulus and peaked at 330 ms, while its polynomial fit peaked at 308 ms.

**Figure 3.**
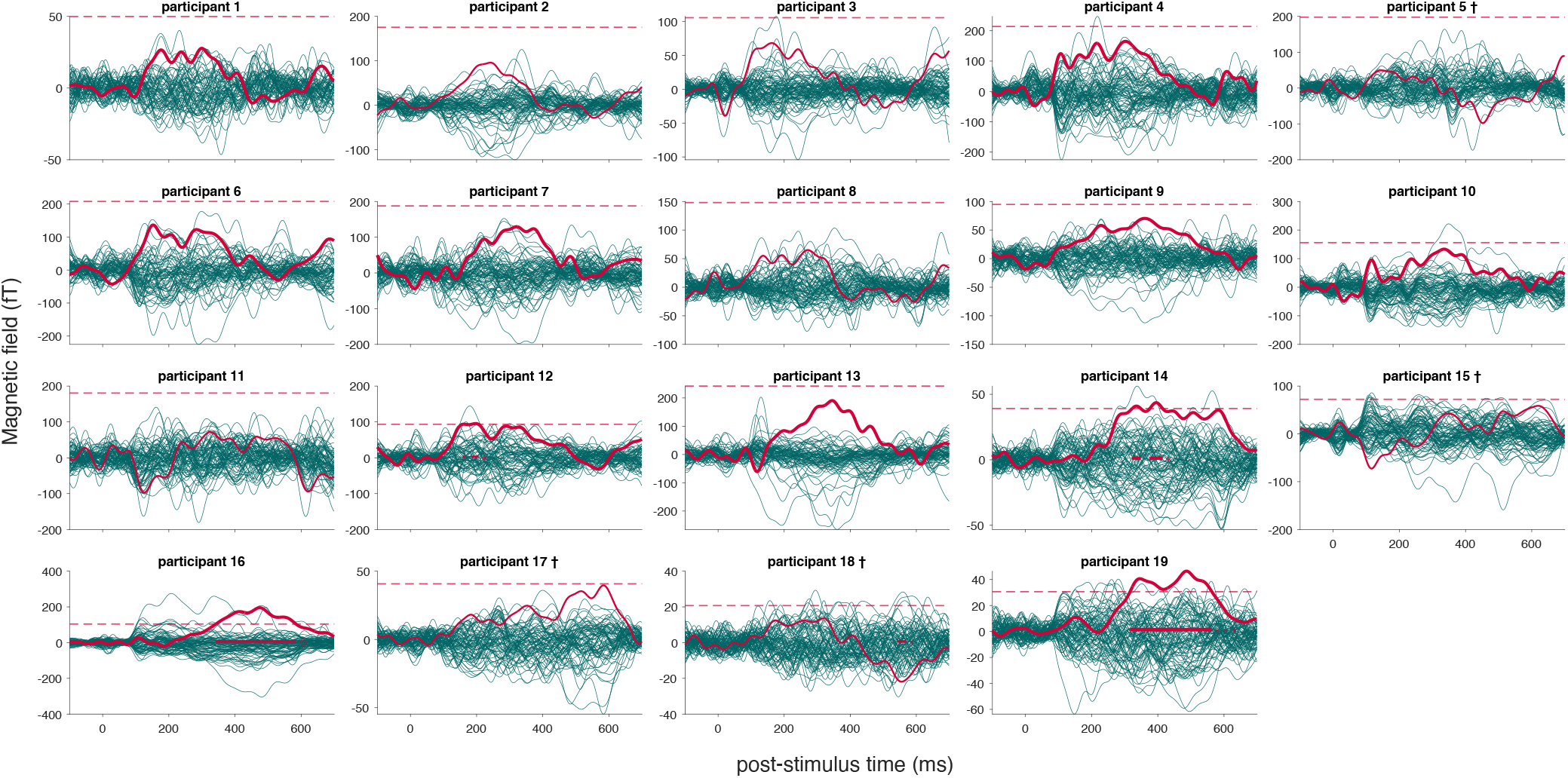
Individual evoked responses (magnetometers, teal) and their PCA1 curves (red) for all fetuses. The PCA1 curves for which the correlation with the group-average response (see Figure 4) was significant are drawn in bold. The dashed lines correspond to the 95^th^ percentile of the surrogate PCA1 curves distribution and the significant periods are highlighted on the time axis. ♢ p<0.0001, *** p<0.001, ** p<0.01 * p<0.05. †Participant excluded from the analysis based on the difference in their response latency compared to the virtual gradiometer response (see Figure S2).

**Figure 4.**
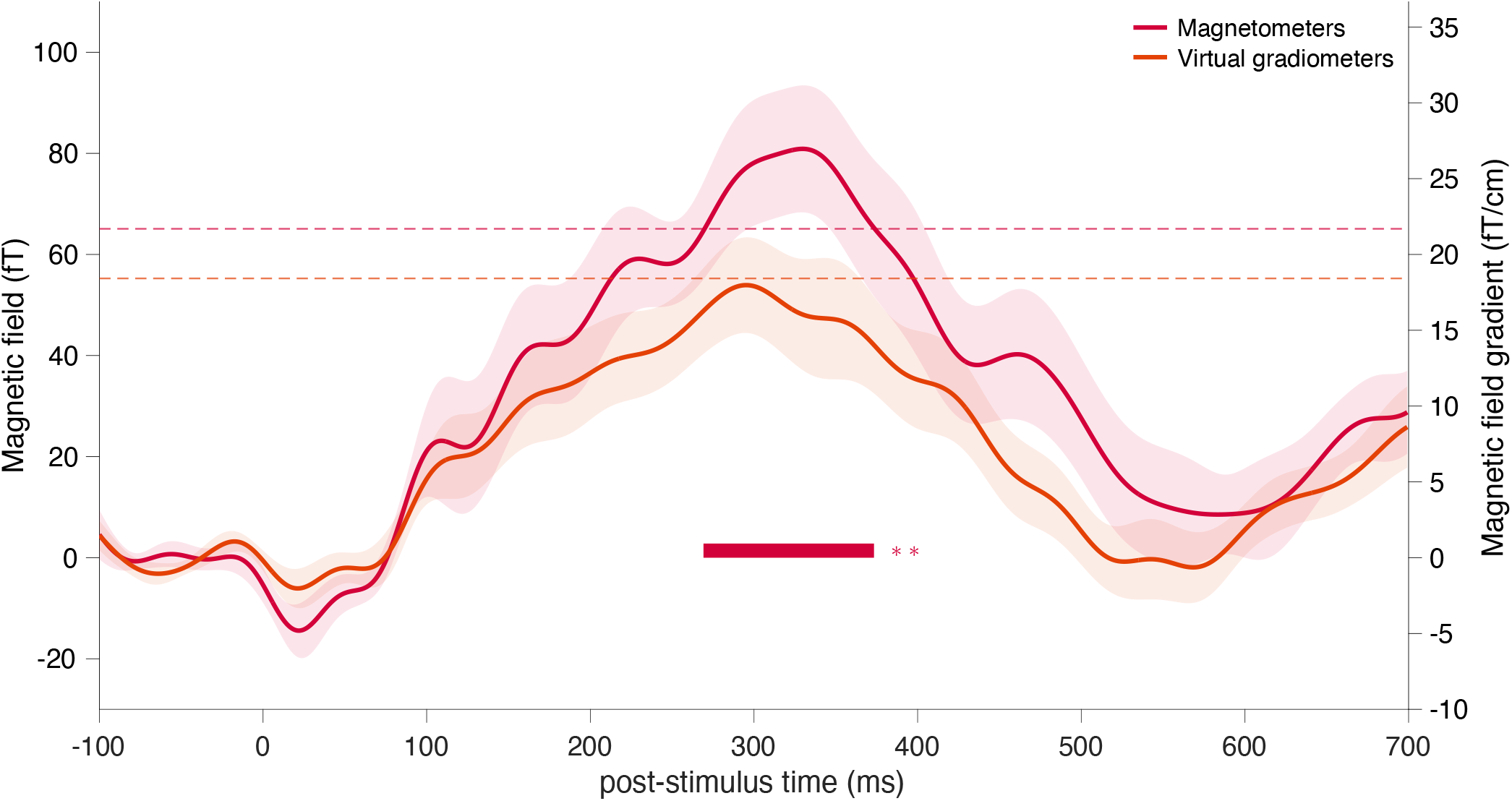
Group average of the PCA1 curves of the evoked responses across fetal subjects, both in terms of magnetometers (red) and virtual gradiometers (orange). Participants with different latencies in magnetometer and virtual gradiometer responses were excluded beforehand. Shaded areas indicate SEM. Statistical tests were performed on these responses (maximum statistics, 10,000 iterations); the significant periods are highlighted on the time axis. **p<0.01. The dashed lines correspond to the 95^th^ percentile of the distributions. The values given for the magnetic field gradient are indicative as they rely on heavy approximations and a 2D custom-made layout.

To further characterize the quality of the individual responses, we computed their lagged correlation with the group-average PCA1 curve (allowing some latency shift) and tested for statistical significance. This lagged correlation was significant in 11 participants, whose PCA1 curves are displayed with a bolder line in Figure 3. All individual amplitudes, latencies, correlations with the group-average PCA1 and the corresponding *p*-values are displayed in Table 1.

**Table 1.** Individual responses characteristics based on PCA1 curves. Amplitude and latency are taken from a polynomial fit of the PCA1 curves. The correlation with the mean is the maximum lagged correlation of the PCA1 curve with the group-average PCA1 curve.d

| Participant | Magnetometers |  |  |  | Virtual Gradiometers |  |  |  |
| --- | --- | --- | --- | --- | --- | --- | --- | --- |
| | Amplitude (fT) | Latency (ms) | Correlation w/ mean | $p$-value of correlation | Amplitude (fT/cm) | Latency (ms) | Correlation w/ mean | $p$-value of correlation |
| 1 | 25 | 254 | 0.83 | 0.038 | 9 | 280 | 0.94 | < 0.0001 |
| 2 | 84 | 234 | 0.84 | 0.049 | 8 | 176 | 0.38 | 0.354 |
| 3 | 65 | 184 | 0.73 | 0.051 | 9 | 206 | 0.81 | 0.037 |
| 4 | 149 | 255 | 0.9 | 0.001 | 30 | 203 | 0.89 | < 0.0001 |
| 5 | 60 | 458 | 0.54 | 0.258 | 29 | 208 | 0.83 | 0.01 |
| 6 | 120 | 227 | 0.86 | < 0.0001 | 44 | 313 | 0.94 | < 0.0001 |
| 7 | 125 | 318 | 0.93 | 0.002 | 27 | 259 | 0.89 | 0.001 |
| 8 | 58 | 218 | 0.75 | 0.089 | 27 | 288 | 0.86 | 0.015 |
| 9 | 60 | 347 | 0.93 | < 0.0001 | 16 | 202 | 0.84 | 0.024 |
| 10 | 109 | 330 | 0.93 | < 0.0001 | 24 | 232 | 0.86 | 0.003 |
| 11 | 60 | 356 | 0.65 | 0.059 | 26 | 196 | 0.77 | 0.022 |
| 12 | 84 | 238 | 0.87 | 0.006 | 26 | 368 | 0.83 | < 0.0001 |
| 13 | 170 | 337 | 0.94 | 0.001 | 37 | 352 | 0.87 | 0.006 |
| 14 | 40 | 383 | 0.92 | < 0.0001 | 9 | 352 | 0.88 | < 0.0001 |
| 15 | 59 | 599 | 0.54 | 0.322 | 20 | 236 | 0.9 | < 0.0001 |
| 16 | 183 | 460 | 0.95 | < 0.0001 | 27 | 465 | 0.84 | < 0.0001 |
| 17 | 36 | 558 | 0.7 | 0.151 | 7 | 304 | 0.89 | < 0.0001 |
| 18 | 18 | 562 | 0.71 | 0.174 | 6 | 347 | 0.9 | < 0.0001 |
| 19 | 43 | 462 | 0.92 | 0.004 | 10 | 422 | 0.91 | < 0.0001 |

#### 3.2.2 Virtual gradiometers

The auditory evoked responses were also evaluated in terms of virtual gradiometers. Individual responses are presented in Figure S2. Their latencies relative to the magnetometer responses were used to determine for which subjects the observed signal did not likely correspond to a neural response. Those subjects were discarded for the group analysis (n=4). For the remaining subjects, the latencies were not significantly different than those obtained from the magnetometer data. The group-average virtual gradiometer response (presented in Figure 4, alongside the previously described magnetometer response) did not reach the significance threshold. Nevertheless, its temporal dynamic does not differ from the magnetometer response. Finally, 18 fetuses had their PCA1 curve significantly correlated with the group-average PCA1 curve.

#### 3.2.3 Comparison with the neonatal responses

The fetal responses described above were compared to the responses observed in newborns. We restricted these analyses to the subgroup of fetuses for which the SOA was comparable to the one used in the neonatal sample, i.e. 500-700 ms (n=13 total, n=11 after discarding participants based on the latency difference between magnetometers and virtual gradiometers). Although the subgroup of fetuses who had longer SOA (700-900 ms, n=6 total, n=3 retained) is too small to make meaningful comparisons, latencies seemed slightly delayed compared to the short SOA group (see Figure S3).

Comparing the latencies observed at each timepoint (fetal and neonatal, see Figure 5a), a significant difference was found for the magnetometer latencies (fetal: mean = 275 ms, SD = 59 ms; neonatal: mean = 238 ms, SD = 42 ms, *p*=0.04, mixed paired/unpaired *t*-test), but not for the virtual gradiometer ones (fetal: mean = 256 ms, SD = 64 ms; neonatal: mean = 224 ms, SD = 52 ms, *p*=0.33). The group average responses per timepoint are presented in Figure 5b.

**Figure 5.**
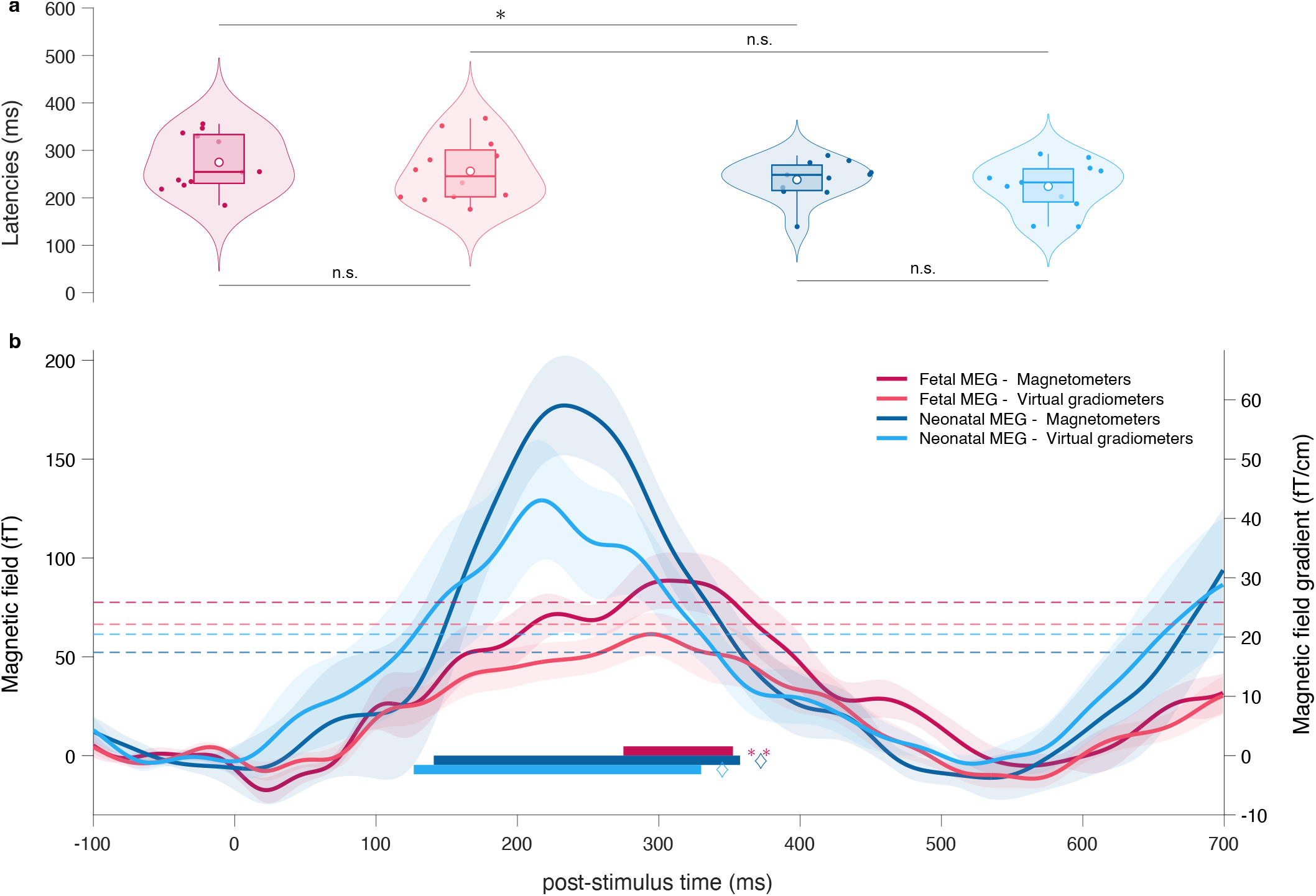
Comparison of the fetal and neonatal responses. (a) Latencies comparison. (b) Group-average PCA1 curves of the evoked responses (magnetometers: dark colors, virtual gradiometers: light colors) for both fetal (red) and neonatal (blue) samples. Shades indicate SEM. Compared to Figure 3, here only the fetuses that had simulations with similar SOA as in the neonatal timepoint were included (n=13 total, n=11 after discarding participants based on the latency difference between magnetometers and virtual gradiometers). All participants of the neonatal timepoint have been included (n=14 total, n=11 retained). Significant periods are highlighted on the time axis. The dashed lines correspond to the 95^th^ percentile of the statistical distributions. ♢ p<0.0001, **p<0.01, *p<0.05, n.s.: non-significant.

For the participants that took part in both timepoints, subject-wise comparisons of their PCA1 curves are displayed in Figure 6.

**Figure 6.**
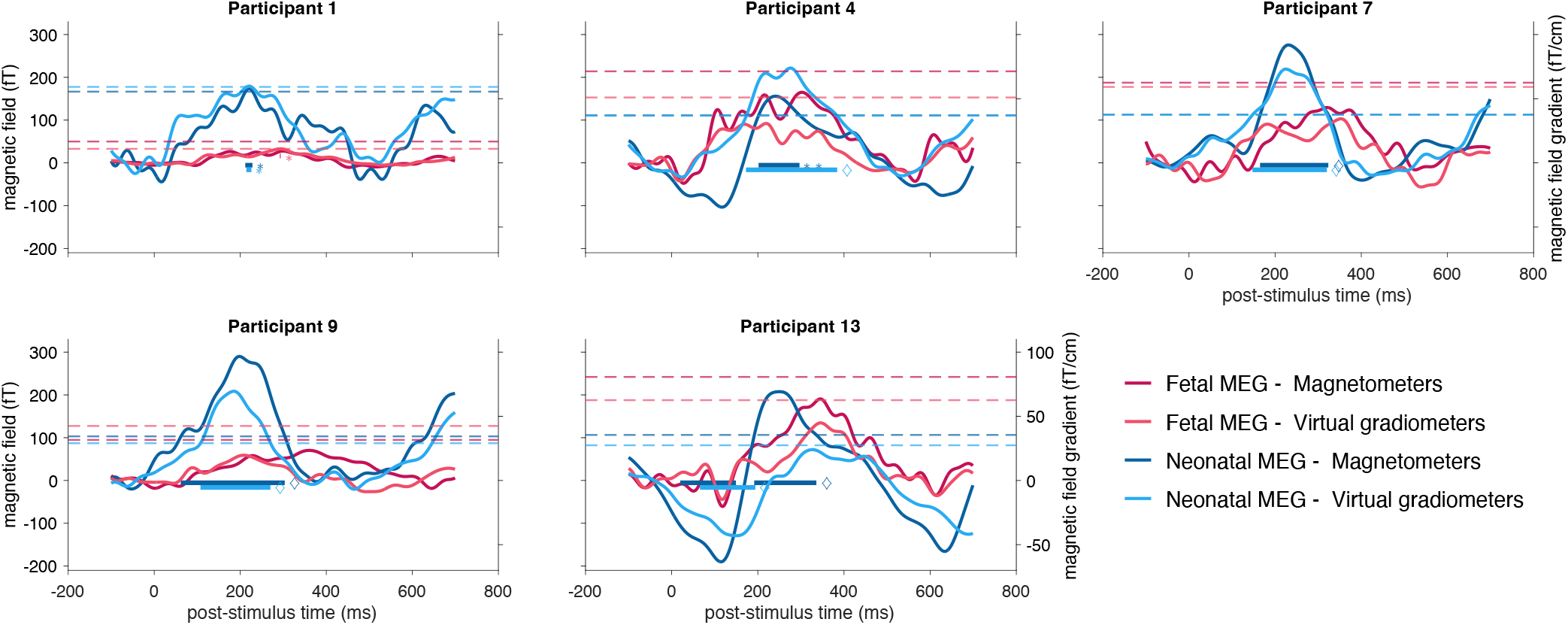
Individual level comparison of the PCA1 curves of the evoked responses in participants that came for both timepoints of this study (n=9) and had similar SOA (n=7 total, n=5 retained). All is as in Figure 5. ♢ p<0.0001, *** p<0.001, ** p<0.01, * p<0.05.

## 4. Discussion

This work provides an unprecedented adaptation of an OPM-MEG system to record fetal brain activity in response to auditory stimulations. This was achieved by designing a comfortable wearable fetal OPM-MEG setup with sensors organized onto an elastic belt that was adjusted over the maternal abdomen. The system was well-tolerated by all pregnant women and allowed to dissipate the heat generated by the OPM sensors.

The first achievement of this work was the possibility to perform fetal MCG for each acquisition. Accordingly, recent works pointed at OPM-MEG as a to-go tool for fetal magnetocardiography (Escalona-Vargas et al., 2024; Strand et al., 2019; Wurm et al., 2023). In the current investigation, fetal and maternal heart traces were easily distinguishable by means of ICA, with fetal participants showing increased heartbeat frequency and heart rate variability throughout the recording compared to their mothers. Mean fetal heart rate data were in line with previous OPM investigations (Escalona-Vargas et al., 2024). Even though MCG was not the main aim of this work, we confirmed the high efficacy of OPM-MEG to perform MCG in utero. These heart rate components, together with other artifacts and sources of noise, were then used to clean the current data and extract neural evoked responses.

The main achievement of the current work is undoubtedly the successful recording of auditory evoked activity in response to 500 Hz tones in a group of fetuses in the third trimester of gestation. Most fetuses had an observable auditory evoked response that peaked in the 200-400 ms time range, even reaching significance at the individual level for a quarter of the participants. However, as fetal head positioning is variable and not always optimal with respect to the sensors array, a response is not expected for all subjects. To better discriminate the fetuses with a plausible brain response, we analyzed the data in terms of virtual gradiometers, which have the advantage of eliminating the homogeneous part of the noise. We discarded for the group analyses the participants for which the magnetometers and virtual gradiometers latencies were too dissimilar.

The group response reached significance and peaked around 300 ms post stimulus exposure. Most retained fetal participants showed an evoked response significantly correlated with the group response (magnetometers: 73%, virtual gradiometers: 93%).

The latency of the evoked response obtained with fetal OPM-MEG is in line with previous cryogenic MEG data, revealing response dynamics in a similar time range (Draganova et al., 2007; Eswaran et al., 2021). Response latencies spanned a relatively wide range across fetal subjects. This is congruent with the classification of early (100 ms to 300 ms) and late responders (300 ms to 500 ms) to standard tones suggested by past fetal research (Draganova et al., 2007). This fetal auditory evoked response has been interpreted as a delayed component compatible with the mature N100 (Preissl et al., 2004). The signal amplitudes found in this study are, for some fetuses, higher than the amplitudes reported in previous cryogenic investigations (often in the 6 to 40 fT range, see (Draganova et al., 2007; Eswaran, Preissl, et al., 2002; Preissl et al., 2001)). However, the factors by which they differ are reasonable and can be explained by the magnetometers being closer to the neural source, i.e., either the OPMs being closer to the fetal brain or the fetuses’ heads being oriented more appropriately, or both. Indeed, the SQUID sensors must be housed in a dewar filled with liquid helium, which increases their distance to the maternal abdomen compared to OPMs, and in turn to the fetal brain.

On top of recording fetal auditory evoked responses using OPM-MEG, an added value of this work is its longitudinal design. In fact, a subset of participants tested in utero came back at 1 month of age to undergo a similar follow-up investigation with on-scalp OPM-MEG. Overall, auditory evoked responses revealed a latency decrease from the fetal to the newborn timepoint. A mixed paired-unpaired comparison of fetal and newborn participants who were exposed to a similar SOA showed a significant latency decrease with age in terms of magnetometers. Nevertheless, the latency difference did not reach significance in terms of virtual gradiometers. Past cryogenic MEG investigations also revealed a latency decrease from fetal to newborn stages, often reaching significance level (Holst et al., 2005; Lengle et al., 2001; Schleussner et al., 2001; Sheridan et al., 2008), and sometimes not (Draganova et al., 2007). While auditory processing capabilities are well-developed during the third trimester of gestation (Iyengar, 2012), the first postnatal weeks can play a critical role in temporal processing of auditory information (Sheridan, Draganova, et al., 2010). A latency decrease can be explained in terms of progressive cortical development and myelination but also increased auditory exposure and experience of the newborn group (Herschkowitz, 1988; Matschke et al., 1994). Interestingly, current data revealed that fetal latencies were much more variable than newborn ones. This could be explained by several factors. First, it could be an intrinsic characteristic of the fetal evoked responses. Overall, considerable intersubject variation has been reported in fetal evoked responses to sounds (Draganova et al., 2007). Second, fetal behavioral states, not recorded in the current study, could be more variable than infant ones. Latency variability of the auditory evoked responses has been linked to the behavioral states, both in newborns (Duclaux et al., 1991; Weitzman et al., 1965) and in fetuses (Kiefer-Schmidt et al., 2013). We also cannot rule out the possibility that the response variability is related to more variable noise levels in the fetal group, which can be due to variable fetal positioning in the womb and distance from the fetus head to the sensors or variable degrees of movement during the recording. On the other hand, brain-sensor distance in the newborn sample was rather fixed.

This work has some limitations that can be addressed in future investigations. First, future research could better investigate the link between fetal sensory processing and individual variables, such as gestational age and maternal factors (which were relatively homogeneous in our sample), and/or behavioral states during the recording. The current study was indeed not designed to investigate fetal behavioral state variations (e.g., awake vs. sleep states) during the OPM-MEG recording. A possible solution to classify fetal behavioral states is measuring fetal heart rate together with fetal movement patterns (Semeia et al., 2022; Vairavan et al., 2016). Other limitations are related to our setup and the associated signal noise levels. In particular, the variable localizations and orientations of our array of OPM sensors precluded the use of advanced denoising approaches based on multipole expansions (Holmes et al., 2023; Tierney et al., 2022, 2024). Respiratory and movement artifacts may also have been associated with our wearable, flexible system. Thankfully, the great adaptability of OPM sensors allows researchers to test different setup configurations. A rigid system would allow for more advanced denoising thanks to the known positions and orientations of the sensors. It would also be less susceptible to noise and artifacts in the recordings because of the limited contact with the participant’s body. However, such a system is less adaptable to different participants, which is a major asset of the current setup. An alternative possibility to decrease artifacts while ensuring a good sensor-brain distance may be a semi-rigid solution with the pregnant participant lying prone. Such bed-based array system has recently revealed successful results for fetal MCG recordings (Escalona-Vargas et al., 2024). Another limitation of the present work is the need of spatial filtering (removal of sound artefact and external noise). This spuriously deforms the topography. Further, topographic maps can get superimposed in case of fetal movements, which could not be tracked during the current data acquisition. Overall, future fetal OPM works could better address noise management and filtering strategies in an environment that is highly affected by noise. OPM-MEG is still an emerging technique, and the quality of fetal recordings will only improve with the future development of new tools, such as the automatic registration of the OPMs localization and orientation using external coils, e.g., the zero field nulling coils of the MSR, or the adaptation to fetal MEG of a halo of fixed head positioning coils (Hill et al., 2024; Livanainen et al., 2022; Pfeiffer et al., 2018, 2020).

Overall, the current work successfully adapted OPM-MEG to study fetal brain responses and paves the way for further investigations on fetal cognition and maturational changes across early development. To date, converging evidence has revealed that the fetus in the third trimester of gestation has already the skills to process sensory stimuli in broadly similar ways to the newborn infant (Dunn & Reid, 2020). While infant cognitive development has been quite extensively studied over the last decades, the study of perception and cognition should start from the fetus (Reid, 2024). In this context, various infant paradigms used to investigate early perceptual and cognitive capabilities could be adapted to fetal samples. The current work demonstrates that OPM-MEG is a suitable and promising tool to study not only fetal cardiac activity but also fetal brain functions. Up to now, fetal SQUID-MEG investigations have been limited to a couple of laboratories worldwide, whereas OPM-MEG, being cheaper and more versatile, has the potential of becoming a large-scale solution. Indeed, fetal SQUID-MEG, such as the SARA system, represents a massive investment only suitable for pregnant participants, while the same OPM-MEG sensors can be adapted to multiple study designs and populations. In the current study, the system has been adapted to both fetal and newborn recordings, and the same OPM system has been successfully used with infants and older children as well as in adults in other studies from our group (Feys, Corvilain, Bertels, et al., 2023; Feys et al., 2022; Feys, Ferez, et al., 2024; Feys, Wens, et al., 2024). This makes OPM-MEG a promising non-invasive and truly lifespan-compliant neuroimaging tool that can be used to test participants of all ages and conditions. In the long run, once fetal neurodevelopment has been well characterized by OPM-MEG, the latter will also have the potential to become a valuable tool in the assessment of at-risk pregnancies.

## Additional Information

### CRediT statement

Conceptualization: JB, XDT

Data Curation: PC

Formal Analysis: PC, VW, CC, JB, XDT

Funding Acquisition: JB, XDT

Investigation: PC, JB, CC, LF, MF, AD, RL, DG, CDC

Methodology: PC, JB, XDT, VW

Project Administration: JB, XDT, PC, CC

Resources: XDT, JB, CC, OF

Software: PC, VW

Supervision: JB, XDT

Visualization: PC

Writing – Original Draft: PC, CC

Writing – Review & Editing: PC, CC, JB, XDT, VW, MF, OF, LF, AD, RL, DG, CDC

### Data and materials availability

All data, code, and materials used in the analyses will be archived to an open data publishing platform and will be made publicly available upon publication.

The consent and authorization to publish the photos have been obtained from the participant (or their legal representative) in Figure 1.

