## Supplemental Material for "Fetal magnetoencephalography based on optically pumped magnetometers"

| Participant | Sex | Gestational age (weeks + days) | Participation in neonatal study | Mean heart rate (bpm) | Heart rate variability (bpm) |
|---|---|---|---|---|---|
| 1 | M | 38 + 6 | X | 127 | 8 |
| 2 | F | 38 + 0 | X | 139 | 1.6 |
| 3 | M | 35 + 3 | | 154 | 10 |
| 4 | F | 38 + 6 | X | 128 | 4 |
| 5 | M | 38 + 4 | X | 130 | 14.2 |
| 6 | F | 37 + 5 | | 136 | 7.1 |
| 7 | F | 38 + 5 | X | 138 | 9.1 |
| 8 | F | 38 + 0 | | 147 | 6.7 |
| 9 | F | 38 + 5 | X | 147 | 12.1 |
| 10 | M | 39 + 4 | | 134 | 6.5 |
| 11 | F | 38 + 4 | | 147 | 4.5 |
| 12 | F | 40 + 1 | X | 143 | 7.2 |
| 13 | M | 39 + 0 | X | 134 | 2.3 |
| 14 | F | 38 + 5 | | 131 | 7.8 |
| 15 | F | 37 + 5 | | 157 | 10.9 |
| 16 | F | 38 + 2 | X | 145 | 8.5 |
| 17 | F | 38 + 4 | | 146 | 9 |
| 18 | M | 38 + 1 | | 132 | 5.7 |
| 19 | F | 37 +3 | | 153 | 10.5 |

***Table S**1. Participants' information and heart rate data.*

| Participant | Recording time (s) | Total length of artifact periods (s) | | Number of stimuli | |
|---|---|---|---|---|---|
| | | Visual inspection | z-score rejection | Presented | Accepted |
| 1 | 425 | 37 | 86 | 759 | 568 |
| 2 | 426 | 8 | 25 | 770 | 706 |
| 3 | 428 | 75 | 133 | 763 | 498 |
| 4 | 429 | 50 | 121 | 766 | 526 |
| 5 | 427 | 78 | 171 | 766 | 427 |
| 6 | 428 | 87 | 160 | 763 | 459 |
| 7 | 418 | 74 | 115 | 756 | 531 |
| 8 | 425 | 77 | 134 | 765 | 496 |
| 9 | 429 | 41 | 93 | 758 | 556 |
| 10 | 422 | 74 | 121 | 754 | 508 |
| 11 | 428 | 49 | 97 | 770 | 581 |
| 12 | 421 | 98 | 153 | 707 | 404 |
| 13 | 428 | 162 | 242 | 768 | 311 |
| 14 | 867 | 121 | 201 | 1051 | 766 |

| | | | | | |
|---|---|---|---|---|---|
| 15 | 428 | 74 | 116 | 526 | 370 |
| 16 | 426 | 122 | 189 | 526 | 285 |
| 17 | 854 | 27 | 124 | 1122 | 939 |
| 18 | 853 | 90 | 169 | 1046 | 802 |
| 19 | 853 | 126 | 226 | 1048 | 723 |

***Table S**2. Recordings information. For each participant, the total recording time, the total length of periods considered as containing artifacts, both after a first visual inspection and a subsequent z-score rejection, and the number of stimuli presented and kept (i.e., not intersecting a period with artifacts) are provided.*

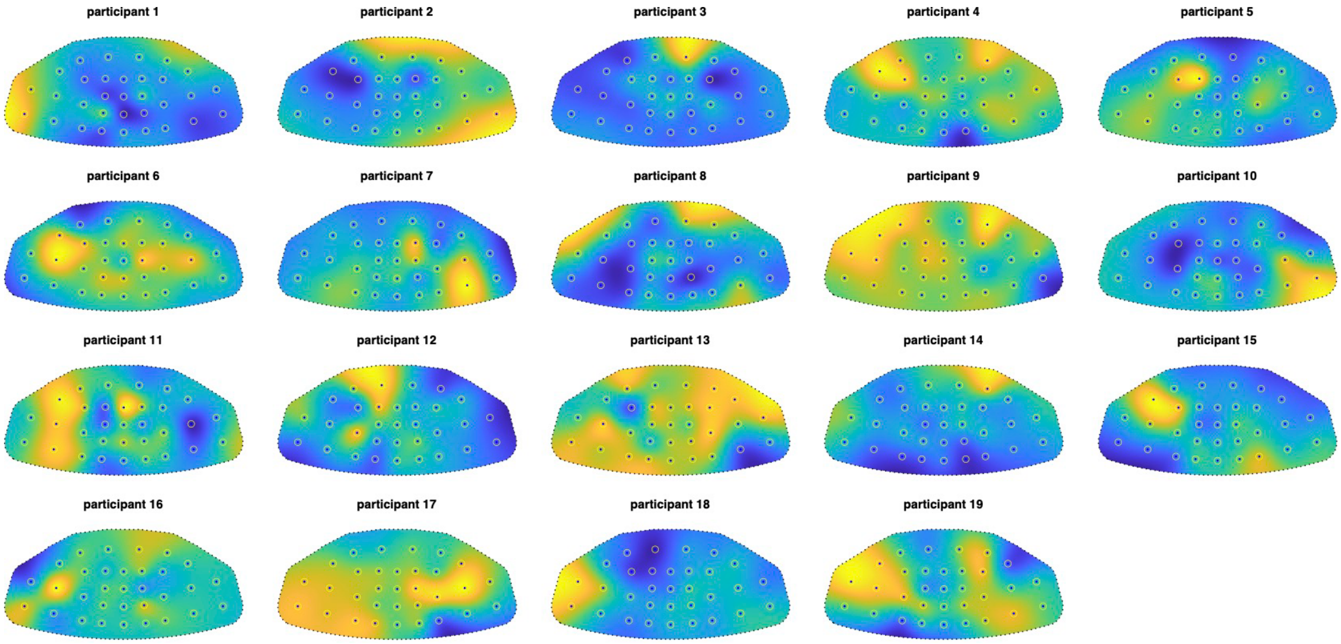


***Figure S1**. Individual radial topographic maps. These maps are computed as the radial part of the spatial map associated with the first principal component of the evoked response. These values thus represent how much each channel contributes to the PCA1, and the actual values are not meaningful, hence we chose not to include scale bars. The 2D layout used for representing the abdomen belt was custom-made.*

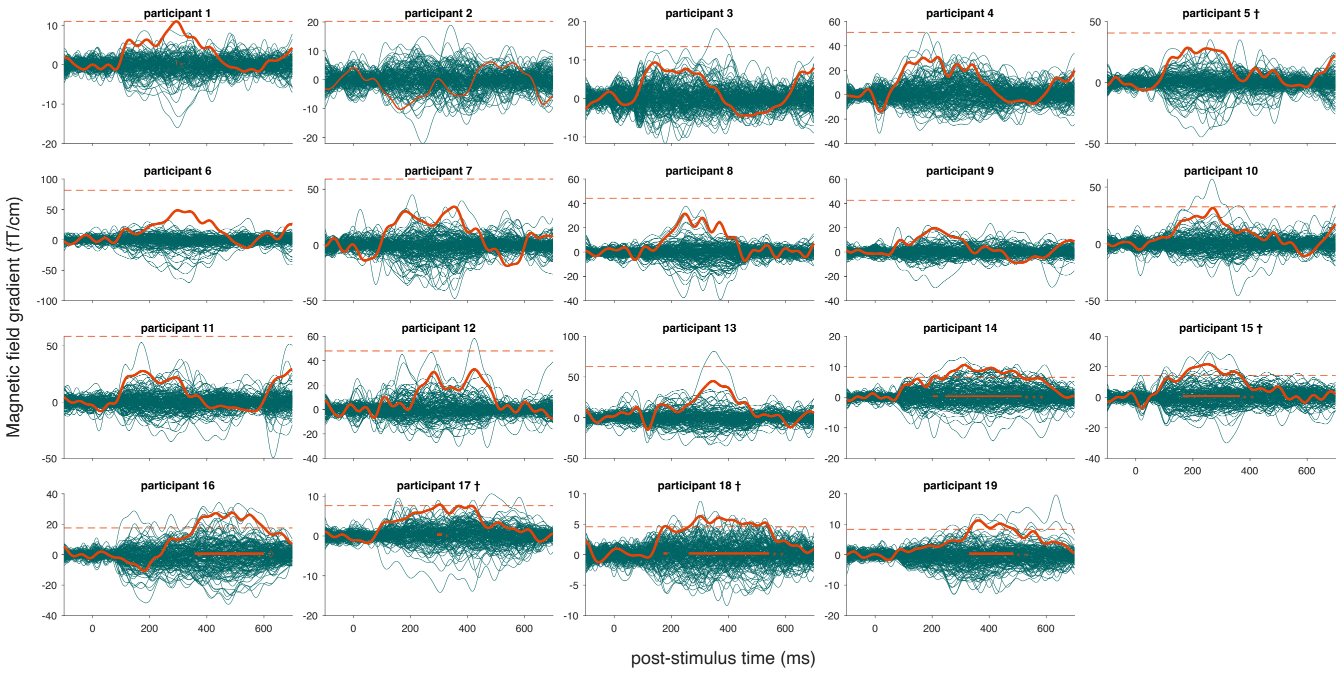


***Figure S2****. Individual evoked responses (virtual gradiometers, teal) and their PCA1 (orange) for all fetuses. The PCA1 curves for which the correlation with the group-average response (see Figure 4) was significant are drawn in bold. The dashed lines correspond to the 95$^{th}$ percentile of the surrogate PCA1 distribution and the significant periods are highlighted on the time axis. ◊ p<0.0001, *** p<0.001, ** p<0.01 * p<0.05. †Participant excluded from the analysis based on the difference in their response latency compared to the magnetometers' response (see Figure 3).*

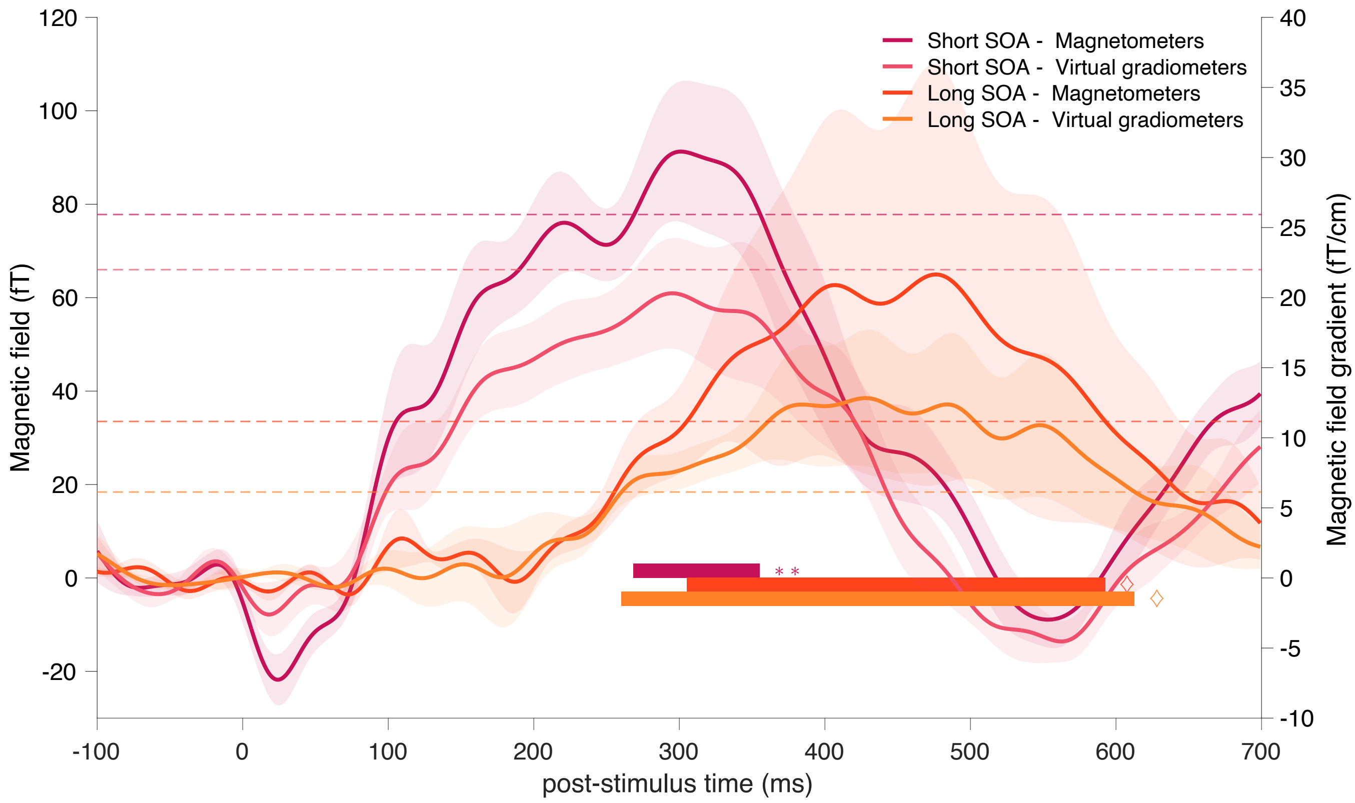


***Figure S3**. Average of the PCA1 curves of the evoked responses across fetal subjects, split into the groups that had shorter (n=13, red) and longer (n=6, orange) sound onset asynchrony, both in terms of magnetometers (darker) and virtual gradiometers (lighter). Statistical tests were performed on this response (maximum statistics, 10000 iterations), the significant periods are highlighted on the x-axis. ◊ p<0.0001, **p<0.01. The dashed line corresponds to the 95<sup>th</sup> percentile of the distributions. The values given for the magnetic field gradient are indicative as they rely on heavy approximations and a 2D custom-made layout.*
